# Extracellular Matrix Stiffness Controls Actin Cytoskeletal Tension

**DOI:** 10.64898/2026.08.02.742292

**Authors:** L. Perego, M. Sergides, C. Arbore, G. Bianchi, L. Gardini, M. Capitanio

## Abstract

Mechanical properties of the extracellular matrix (ECM) regulate cell behavior and strongly impact the cytoskeleton, yet direct measurements of actin tension across different ECM stiffnesses are lacking. Using a genetically encoded FRET-based actin tension sensor, we show that cells cultured on stiffer substrates exhibit increased spreading, reduced multicellular aggregation, and higher molecular tension within filamentous actin. Complementary measurements using an α-actinin tension sensor support increased cytoskeletal mechanical loading. These results provide direct evidence that ECM stiffness modulates intracellular actin tension during mechanotransduction.

## Main text

Cells continuously sense and adapt to the mechanical properties of the extracellular environment, which regulate cell shape, adhesion, actomyosin contractility, and motility. On the other hand, aberrant mechanical cues contribute to pathologies such as cancer progression and metastasis. In particular, substrate stiffness strongly influences cell morphology, with increased stiffness associated with enhanced cell spreading on the adhesive surface^1^. These morphological changes are accompanied by increased focal adhesion maturation and density, as well as by higher traction forces exerted by cells^2,3^. Moreover, increased substrate stiffness induces reorganization of the actin cytoskeleton, characterized by enhanced stress fiber formation, increased actin density, and a global increase in cell stiffness^1,4^. Despite the strong interplay between extracellular mechanics and the cytoskeleton, direct measurements of how actin cytoskeletal tension responds to physiological tissue stiffnesses are still lacking. Addressing this question is essential for understanding how mechanical signals propagate within cells and thereby regulate cell physiology, yet it is hindered by the technical challenges associated with measuring intracellular forces.

Molecular force sensors are invaluable tools for measuring intracellular forces in living cells. Guo et al. developed AcpA (actin-cpstFRET-actin), a genetically encoded force sensor that incorporates the cpstFRET module, a circularly permuted fluorescent protein FRET pair sensitive to mechanical load^5^, between two β-actin monomers^6^. This design enables the sensor to incorporate into filamentous actin (F-actin) and report changes in molecular tension through variations in FRET efficiency (Fig. 1a). Its localization to F-actin and faithful incorporation into the actin cytoskeleton were previously validated by colocalization with F-actin staining and the responsiveness of AcpA to changes in cytoskeletal tension has been extensively validated^6^.

**Figure 1.**
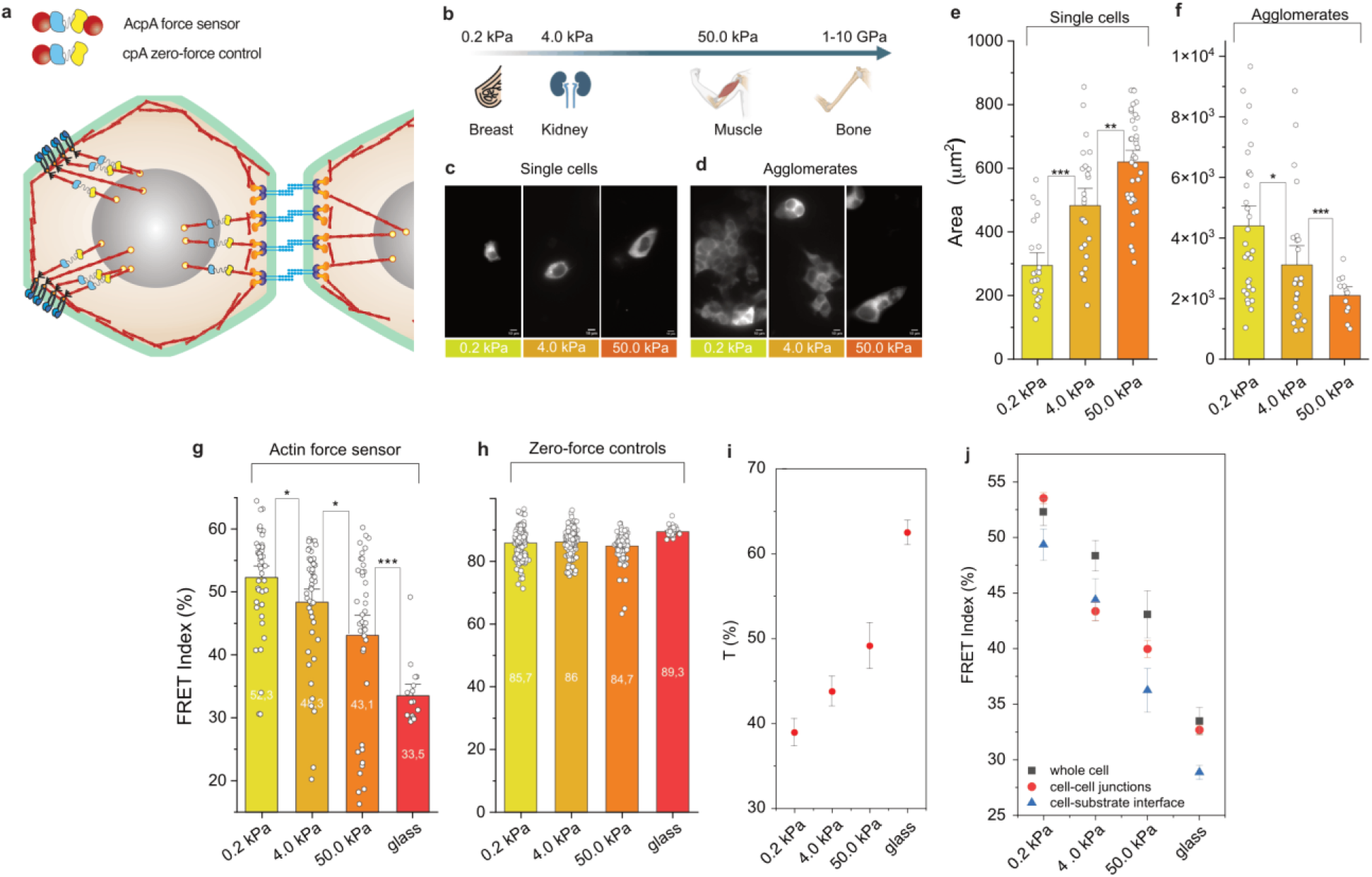
**a)** Schematic of the AcpA force sensor, the cpA zero-force control, and how the sensor is incorporated into F-actin in cells. **b)** Hydrogel stiffnesses used in this study, mimicking breast (0.2kPa), kidney (4.0 kPa), and muscle (50.0 kPa) tissues, and glass (1-10 GPa), mimicking bones. **c**,**d)** Representative fluorescence images illustrating greater spreading of single cells on stiffer substrates (c) and larger cell agglomerates on soft substrates (d). **e)** Single cell area on substrates of increasing stiffness (n = 22, 25, and 38 cells for 0.2, 4, and 50 kPa, respectively). **f)** Aggregate area showing increased clustering on soft substrates (n = 30, 25, and 12 cells for 0.2, 4, and 50 kPa, respectively). **g)** FRET index of the actin tension sensor (AcpA) decreases with increasing stiffness (n = 44, 46, 39, and 17 cells for 0.2, 4, 50 kPa and glass, respectively). **h)** FRET index of zero-force control sensor (cpA) shows no stiffness dependence (n = 228, 198, 139, and 39 cells for 0.2, 4, 50 kPa and glass, respectively). p<0.0001 betwen AcpA and cpA for each condition (0.2, 4, 50 kPa and glass). **i)** The tension index reports increasing actin mechanical loading with substrate stiffness. **j)** Spatial analysis of AcpA FRET index for the whole cell, cell-cell junctional regions, and cell-substrate interface. Bars represent mean ± SE. Statistical significance was assessed using two-tailed Student’s t-test. ns, P *≥* 0.05; *P < 0.05; **P < 0.01; ***P < 0.001; ****P < 0.0001.

Other force sensors that measure F-actin-related forces are based on actin binding proteins, such as α-actinin^6,7^, which embeds into stress fibers, Filamin-A, an actin-binding protein that crosslinks actin filaments within the cortex and stress fibers^8^, or a recently developed force sensor based on two F-tractin binding domains, which act like an actin cross-linking protein^9^. Differently from AcpA, α-actinin, Filamin-A, and F-tractin based force sensors report forces that promote the relative translocation of actin filaments, rather than forces within individual filaments.

To our knowledge, the only previous direct study of the effect of substrate stiffness on actin tension used the AcpA sensor to compare the long-term response (2–7 days) of cells cultured on PDMS and glass substrates^6^. That study reported higher actin tension on the softer PDMS substrate (approximately on the order of hundreds of kPa), which was interpreted as a consequence of cell reprogramming and global cytoskeletal remodeling. By contrast, direct measurements of the short-term response of actin tension to calibrated substrates spanning the physiological range of tissue stiffnesses are still lacking.

To address this, we employed AcpA to report molecular tension within F-actin in HEK cell lines stably expressing AcpA. Cells were cultured on hydrogels with stiffnesses of 0.2, 4.0 and 50.0 kPa, covering a physiological stifness range of soft tissues, as well as on glass coverslips (~1GPa), representing a non-physiological stiff reference (see Methods for more details). These stiffnesses mimic the elastic moduli of breast, kidney, and muscle, respectively, with glass approximating the stiffness of bones^10^ (Fig. 1b). We used cpA (cpstFRET-actin), in which the cpstFRET module is fused to a single actin monomer, as a zero-force control sensor preventing force transmission when incorporated into F-actin (Fig. 1a)^6^.

Consistent with previous observations, cell morphology strongly depended on substrate stiffness^11^. On soft (0.2 kPa) hydrogels, cells formed large multicellular aggregates and adopted rounded shapes, whereas on stiffer substrates (4-50 kPa) cells spread extensively and formed smaller clusters (Fig. 1c,d). Quantification of single-cell area confirmed a marked increase with substrate rigidity (Fig. 1e). Aggregates were significantly larger on soft hydrogels, as demonstrated both qualitatively (Fig. 1d) and quantitatively (Fig. 1f). This behavior is consistent with reduced extracellular matrix (ECM) adhesions and increased cell-cell junctions.

We then analyzed FRET efficiency using a calibrated ratiometric FRET index (see Sergides et al.^12^ and Methods for details) to compare tension levels across conditions. Zero-force cpA controls showed no significant differences in FRET efficiency across substrate stiffnesses, as expected for a sensor that is not mechanically loaded (Fig. 1h). In striking contrast, cells expressing the AcpA sensor exhibited a progressive decrease in FRET efficiency with increasing substrate stiffness (Fig. 1g and Supplementary Fig. S1). Because lower FRET efficiency corresponds to a greater force, these results directly demonstrate an increase in mechanical loading within F-actin as cells adhere to stiffer matrixes. Plot of the tension index T, defined as the normalized FRET index difference between zero-force control and sensor (see Methods), directly highlights the increase in actin mechanical loading with substrate stiffness (Fig.1i).

FRET measurements were acquired by focusing on the adhesion surface; therefore, they reflect the average molecular actin tension within basal cortical actin. To better separate the contribution of actin tension between cortical actin, actin at the cell-substrate adhesion and actin at cell-cell junctions, we produced separated spatial analysis of FRET values. Fig. 1j shows that in all regions, the FRET signal decreases with substrate stiffness, indicating that the tension on the actin cytoskeleton increase globally in the cell. Interestingly, FRET at cell-substrate interface is lower than in other regions and, thus, these regions are where most tension accumulates on the actin cytoskeleton. On the other hand, actin tension at cell-cell junctions is higher than the average cell value in the 4-50 kPa range.

To decouple the effect of the biochemical composition of the adhesion substrate from that of matrix stiffness on actin tension, we compared FRET indexes of cells cultured on bare glass versus glass coated with fibronectin or poly-L-lysine. Despite measurable morphological changes, no significant differences in FRET index were observed among the three conditions, indicating that the changes in FRET index can be attributed specifically to substrate stiffness (see Supplementary Fig. S2).

Finally, we investigated tension in α-actinin, an actin-binding protein that forms antiparallel homodimers, cross-links actin filaments, and connects F-actin to several adhesion proteins. To this end, we employed an α-actinin tension sensor based on the same cpstFRET sensor module^6^ (Fig. 2a). Because α-actinin localizes predominantly to stress fibers, focal adhesions, and cell–cell junctions, it provides complementary, yet distinct, information from the actin sensor about the mechanical state of actin-associated structures and adhesion sites. HEK cell lines stably expressing α-actinin-cpstFRET were cultured on hydrogels with stiffnesses of 0.2, 4.0, 50.0 kPa, and on glass coverslips, as before. Again, single-cell area markedly increased with substrate rigidity (Fig. 2b), while cell aggregates were significantly larger on soft hydrogels (Fig. 2c). As previously reported^1^, cells cultured on soft substrates did not exhibit prominent stress fibers and showed low α-actinin fluorescence signals. FRET analysis was restricted to regions near cell–cell adhesion sites within multicellular aggregates, where we observed high α-actinin fluorescence signals. Lower FRET indexes were observed with increasing substrate stiffness (Fig. 2d and Supplementary Fig. S3), while zero-force controls showed no significant differences in FRET efficiency between 0.2 and 4 kPa (Fig. 2e), indicating higher tension in α-actinin near cell-cell junctions. FRET indexes at higher substrate stiffnesses (50 kPa and glass) cannot be directly compared because of a drop in the control FRET. Interestingly, analysis of the tension index indicates that α-actinin tension saturates at these higher substrate stiffnesses (Fig. 2f).

**Figure 2.**
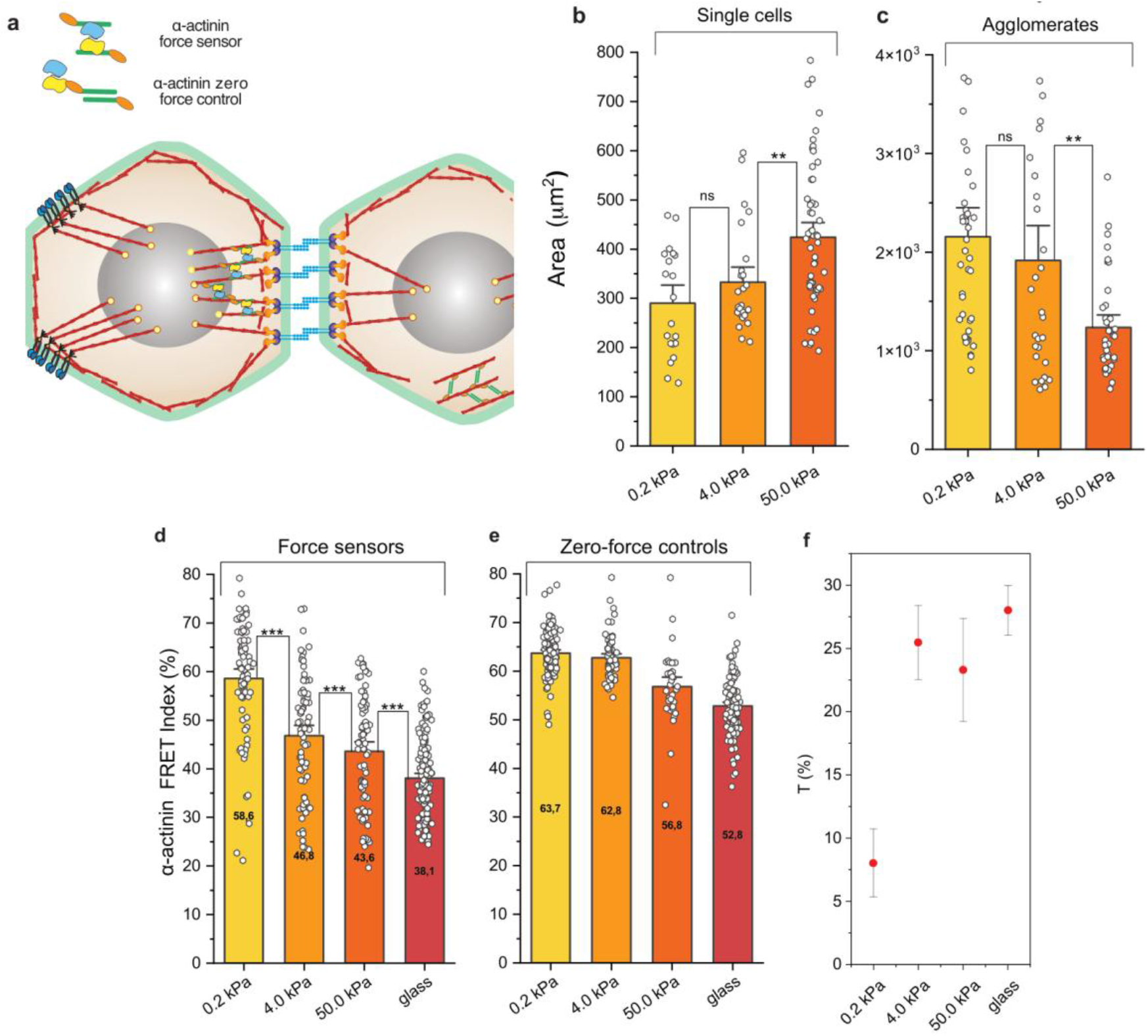
**a)** Schematic of the α-actinin FRET-force sensor, and how the sensor is incorporated into F-actin cytoskeleton in cells. **b)** Single cell areas on substrates of increasing stiffness (n = 20, 26, and 54 cells for 0.2, 4, and 50 kPa, respectively). **c)** Aggregate area showing increased clustering on soft substrates (n = 41, 29, and 37 cells for 0.2, 4, and 50 kPa, respectively). **d)** FRET index of the α-actinin tension sensor versus substrate stiffness (n = 79, 74, 76, and 140 cells for 0.2, 4, 50 kPa and glass, respectively). **e)** FRET index of zero-force control sensors versus substrate stiffness (n = 119, 68, 35, and 123 cells for 0.2, 4, 50 kPa and glass, respectively). p<0.0001 betwen AcpA and cpA for each condition (0.2, 4, 50 kPa and glass). **f)** The tension index reports lower α-actin mechanical load at low substrate stiffness. Bars represent mean ± SE. Statistical significance was assessed using two-tailed Student’s t-test. ns, P *≥* 0.05; *P < 0.05; **P < 0.01; ***P < 0.001; ****P < 0.0001.

Together, these results provide, to our knowledge, the first direct intracellular measurements of how substrate stiffness modulates actin mechanical loading, supporting a role for actin filaments as tension sensors that convert substrate rigidity into cellular responses. Previous measurements of actin tension using the same AcpA sensor in cells cultured for several days on relatively soft PDMS substrates surprisingly reported higher forces than in cells grown on glass^6^. This effect was attributed to cell re-differentiation toward a stem-like state. In the present study, cells were allowed to adhere to substrates for a short period (16-24h), to highlight the effect of cell adhesion rather than long-term cell reprogramming. Our data indicate that, on short timescales, adhesion to soft substrates leads to a global decrease in actin cytoskeletal tension, within both the basal actin network and actin near cell-cell junctions. This observation provides direct evidence that the previously reported reduction in cell traction forces and focal adhesion maturation on soft substrates^2,3^ is accompanied by a reduction in the molecular loading of actin filaments, including at cell-cell junctions, thereby supporting an actin-mediated mechanism linking ECM stiffness to the regulation of cell-cell adhesion^13^. It should be noted that the term *molecular tension*, used throughout this article, is a simplification of the mechanical quantity reported by FRET-based force sensors. These sensors are not exclusively sensitive to tensile forces, but may also be influenced by compression, bending, torsion, as well as by conformational changes within the sensor itself, including the recently reported mechanical switching of fluorescent proteins^9^. Despite these limitations, which preclude a direct quantitative conversion of the measured FRET index into force values (see Supplementary Information), our conclusions are based on relative comparisons of the FRET index measured with the same sensor under different experimental conditions. Such comparisons remain indicative of changes in the mechanical loading experienced by the actin cytoskeleton and by α-actinin.

Our results highlight that increases in substrate stiffness rapidly propagates into higher actin tension. Interestingly, the AcpA FRET index was similar in cells cultured on fibronectin- and poly-L-lysine-coated substrates, despite measurable differences in cell morphology and the established impact of these substrates on focal adhesion maturation and traction force^3,14^. One possible explanation is that changes in integrin-mediated adhesion and total traction forces do not necessarily result in proportional changes in the mechanical loading experienced by individual actin filaments, which is the quantity reported by AcpA. Although the molecular basis of this observation remains to be established, it suggests that the substrate stiffness-dependent increase in actin mechanical loading observed here cannot be explained solely by differences in adhesion biochemistry.

F-actin has been proposed to function as a tension sensor, where tension on actin modulates the binding of several actin-binding proteins, including myosin. Moreover, increased tension within the actin cytoskeleton has been suggested to propagate from cell-ECM and cell-cell linkages toward the nucleus, thereby influencing gene expression and cell fate^15^. By directly measuring actin tension across substrate stiffnesses, our work supports a mechanotransduction model in which ECM mechanics is processed and propagated into cells as changes in F-actin tension, which might modulate intracellular biochemical processes and regulate gene expression. Our results offer a methodology and a quantitative basis for future investigations into how mechanical cues are transmitted from the extracellular environment to intracellular processes.

## Methods

### Sample handling and preparation

HEK293 cells stably expressing actin- or α-actinin-based FRET force sensors^6^ were thawed from −80°C aliquots under sterile conditions. Each 1 ml frozen vial was diluted in 4 ml pre-warmed (37°C) culture medium, centrifuged (3 min, 200x g) to remove the DMSO-containing freezing medium, and resuspended in fresh medium before plating in 60 mm dishes. Cells were maintained in a CO_2_ incubator and passaged at least twice prior to imaging.

Cells were plated on either glass or EasyCoat™ Softslip hydrogels (Matrigen) with varying, precalibrated stiffness values. EasyCoat™ Softslip hydrogels are synthetic polyacrylamide gels crosslinked with bisacrylamide. Differences in stiffness are achieved by varying the percentage of bisacrylamide, while keeping the polyacrylamide concentration constant. The EasyCoat™ gels are functionalized with quinone groups, which form covalent bonds with molecules containing primary amines, thiols, or other strong nucleophiles. In our experiments, the hydrogels were pre-coated with fibronectin to enhance cell adhesion.

For experiments on glass, cells were plated at ~10% confluency on 18 mm glass coverslips in 12-well plates. For experiments on substrates of varying stiffness, cells were plated at ~15% confluency on EasyCoat™ Softslip hydrogels (Matrigen). After 16-24h, samples were washed with PBS and mounted in imaging chambers containing Leibovitz’s medium. For hydrogels samples, custom-made PDMS chambers were used (Supplementary Fig. S4). Briefly, a microscope glass slide was coated with a thin PDMS layer, and a PDMS frame of adjustable height (depending on sample thickness) was attached on top. The sample was then inserted into the frame, and the remaining volume was filled with imaging buffer. The chamber was sealed with a glass coverslip and turned upside-down on the microscope stage for imaging with illumination from below.

### Image acquisition

Prior to FRET acquisition, a control sample of HEK cells lacking FRET sensors was imaged under both 445 nm and 514 nm excitation to assess autofluorescence. All FRET measurements were performed on living cells after 16–24 h of adhesion to the substrates, thereby reporting the steady-state molecular tension within the actin cytoskeleton.

Glass samples were imaged in total internal reflection fluorescence (TIRF) configuration on a custom-built setup^12^, whereas samples of cells grown on hydrogels were imaged in highly inclined (HILO) configuration^16^, focusing on the basal adhesion plane. Donor and FRET channels were acquired with 445 nm excitation (50 ms exposure time; 2.5 mW laser power), and the acceptor channel with 514 nm excitation (50 ms exposure time; 2.5 mW laser power). Bright-field images of each field of view were also collected. All images were subsequently processed in MATLAB, for channel alignment, and ImageJ, to assess FRET efficiency, as described below.

### FRET and tension analysis

The images from the three acquisition channels were analyzed using the ImageJ plugin PixFRET, which generates two output images: (i) a FRET image, in which a pixel-wise FRET index is calculated and displayed using a color map, and (ii) a normalized FRET (NFRET) image, in which the FRET index is normalized to obtain a ratiometric FRET value independent of fluorophore concentration. The normalized FRET index (NFRET) was calculated as:

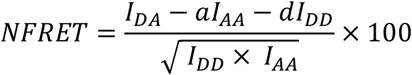

where *I*_*DA*_ is the fluorescence intensity measured in the acceptor channel upon donor excitation, *I*_*AA*_ is the fluorescence intensity measured in the acceptor channel upon acceptor excitation, and *I*_*DD*_ is the fluorescence intensity measured in the donor channel upon donor excitation. The coefficients *a* and *d* represent the spectral bleed-through ratios, accounting for the fraction of fluorescence signal detected in the FRET channel that does not arise from energy transfer.

The NFRET index was previously calibrated using samples with calibrated FRET efficiency values^12^. Therefore, we could estimate FRET efficiency from NFRET. Moreover, the cpstFRET sensor module was previously calibrated using DNA springs^5^. By combining these two calibrations, we could estimate molecular tension from NFRET values (see Supplementary Information).

The tension index T is defined as the percentage difference between the FRET index of the zero-force control and that of the tension sensor, normalized to the zero-force control value:

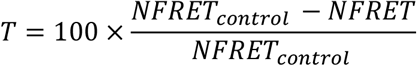

Since the zero-force FRET efficiency can vary between samples due to environmental factors, the T index rescales differences in FRET efficiency between the sensor and the control relative to the dynamic range under the specific experimental conditions.

## Acknowledgments

Funded by European Union - Next Generation Eu, PRIN 2022, MUR 2022T9RM8A. Work supported by #NEXTGENERATIONEU (NGEU) and funded by the Ministry of University and Research (MUR), National Recovery and Resilience Plan (NRRP), project MNESYS (PE0000006) (DN. 1553 11.10.2022). Co-funded by the European Union under HEU-GA 101131771 Lasers4EU.

## Authors’ contribution

L.P. performed experiments and analyzed data, M.S. analyzed data, C.A. and G.B. prepared samples, L.G. supervised experiments, M.C. conceived and supervised experiments. L.P. and M.C. wrote the manuscript. All authors reviewed the manuscript.

